# BURST enables mathematically optimal short-read alignment for big data

**DOI:** 10.1101/2020.09.08.287128

**Authors:** Gabriel Al-Ghalith, Dan Knights

## Abstract

One of the fundamental tasks in analyzing next-generation sequencing data is genome database search, in which DNA sequences are compared to known reference genomes for identification or annotation. Although algorithms exist for optimal database search with perfect sensitivity and specificity, these have largely been abandoned for next-generation sequencing (NGS) data in favor of faster heuristic algorithms that sacrifice alignment quality. Virtually all DNA alignment tools that are commonly used in genomic and metagenomic database search use approximate methods that sometimes report the wrong match, and sometimes fail to find a valid match when present. Here we introduce BURST, a high-throughput DNA short-read aligner that uses several new synergistic optimizations to enable provably optimal alignment in NGS datasets. BURST finds all equally good matches in the database above a specified identity threshold and can either report all of them, pick the most likely among tied matches, or provide lowest-common-ancestor taxonomic annotation among tied matches. BURST can align, disambiguate, and assign taxonomy at a rate of 1,000,000 query sequences per minute against the RefSeq v82 representative prokaryotic genome database (5,500 microbial genomes, 19GB) at 98% identity on a 32-core computer, representing a speedup of up to 20,000-fold over current optimal gapped alignment techniques. This may have broader implications for clinical applications, strain tracking, and other situations where fast, exact, extremely sensitive alignment is desired.

## Introduction

As the amount of next-generation DNA sequencing data increases at a higher rate than computational power(1), approximate heuristic solutions to the fundamental DNA alignment/mapping problem are increasingly used(2). Paradoxically, it seems, the more data made available through advances in sequencing throughput, the less accurate the alignment algorithms used to analyze it. Algorithms with maximal theoretical sensitivity and specificity under mismatch constraints exist but have largely been avoid for high-throughput alignment tasks on next-generation sequencing data in favor of approximate techniques providing faster alignment rate. However, using approximate database search techniques decreases absolute alignment quality in terms of precision and recall.

The problem of short read mapping can be recast as one of fuzzy database search. A database is formed from numerous nucleotide sequences about which some information is known (the so-called “references”); using this database, lookups are then performed for shorter substrings about which little or no information is known (“queries”). A successful lookup assigns meaning to the query, be it match identity, relative position, functional content, taxonomy, or other information as conferred though association with the matched reference(s). These assigned meanings themselves are then treated, often in aggregate, with statistical theory to arrive at some biological insight.

The role of the lookup function in this fuzzy database search problem is simply to provide meaningful mappings from queries to references. As such, the precise technical underpinnings are arguably irrelevant in cases where any such mapping, regardless of the potential existence of closer matches or the number and distribution of other valid matches, provides sufficient information to annotate queries with the desired meaning. Thus, the simpler the information required of a match, the simpler the mapping function can be to provide adequate information. A trivial example is the taxonomic assignment of a single-organism sample to one of two highly divergent taxonomies; an approximate match of a query to any genomic information in one or the other suffices to reasonably infer the taxonomic origin of that query, and the sample’s final assignment can be inferred by tallying the votes of all queries in the sample.

The converse also holds, in that when more complex information is desired, the mapping function must likewise increase in sophistication or accuracy. For instance, if information such as the uniqueness of a match is required, or if the best match is required among many similar matches to resolve minute differences, a lookup function sufficiently sensitive to consider and discriminate among many close alignments to select the correct match will be preferable to one that would merely select a close match.

The technical underpinnings of the mapping function determine how finely it can discriminate between potential database matches. Examples of simple mapping functions are those using exact match (k-mers, substring search) and other alignment-free approaches; the most complex mapping functions involve full gapped alignment often implemented via dynamic programming(3,4). This technique is the foundation for optimal gapped alignment, as well as other fuzzy matching techniques such as bitap(5) or edit distance kernels. Hybrid functions include popular “seed-and-extend” heuristics(6) which combine a simple initial search (the seed) with dynamic programming (the extension) to produce mappings of intermediate stringency and quality.

### Previous work

Efforts to accelerate optimal alignment have largely focused on using efficient data structures(7) and taking advantage of low-level hardware capabilities(8). Some efforts have also focused on pre-processing a database of reference sequences(9,10). To our knowledge, however, there have been no previous efforts to bring optimal short-read mapping to next-generation-scale datasets with the ability to elucidate all ties and perform interpolation to disambiguate tied matches deterministically.

In comparison to general-case alignment, short-read alignment, particularly of the metagenomic variety, presents distinct challenges(11). Perhaps the most obvious among these challenges is the requirement to map millions or billions of short DNA sequences to databases containing thousands of organisms with various degrees of interrelatedness. Less obvious is the need for meaningful disambiguation when a read maps to numerous genomes or chromosomal locations, or the need to aggregate reads into taxonomic bins with a certain level of confidence (the “taxonomic binning problem” in metagenomics). The size of the databases required is large, often containing numerous redundancies (tandem repeats, multiple genomes from a single bacterial species, etc.). The theoretical number of alignments required to perform an exhaustive search is staggering even at modest scales(12); naively aligning one million short reads against a database of one million genes would require one trillion alignments, each of which necessitating a number of calculations proportional to the product of the lengths of query and reference.

However, these challenges of short-read alignment are not without attendant opportunities for substantial performance optimization. There are numerous restrictions embedded in the problem definition which afford opportunities to reduce computational complexity. Since the alignments are typically DNA (or reverse-transcribed RNA), the effective alphabet size is small (usually 4 letters, but up to 16 with IUPAC ambiguity codes(13)). Since “mapping” implies the presence of close matches in the database, tools for exploring more remote evolutionary homologies (local alignment, affine gaps, and evolutionary scoring matrices) may be excluded for certain purposes. Indeed, allowing for some degree of mismatch, each query sequence is expected to be fully contained within some reference in order for a mapping to qualify. Further, particularly in metagenomics applications, a uniform selection criterion such as sequence identity (BLAST(14) percent id), or some number of mismatches, is often imposed by which to filter out potentially spurious alignments. Such a criterion also naturally gives rise to a simplified scoring framework by which to compare alignments across queries and references, and such a scoring scheme (percent identity) is used in the literature to define taxonomic homology thresholds for marker genes.

There are other restrictions that can be imposed on the mapping problem to improve computational efficiency. Databases are relatively immutable; that is, it is reasonable to expect that multiple query datasets will be aligned against a single reference database, allowing pre-processing or indexing of this database to facilitate future alignments. Because of the nature of short-read datasets, query sequences produced by a given sequencing technology have a well-defined maximum length. This is particularly applicable when the database contains numerous repeats, as this maximum length can define a “duplication window” with which to abstract away duplicated portions of reference sequences during database pre-processing, espousing concepts from compressive genomics and allowing database size to scale sub-linearly with the number of reference sequences therein. In addition to redundancy in the database, we have observed that by sorting the query sequences alphabetically we can use the high level of redundancy in the prefix of one query sequence and the next query sequence to reduce search time.

Further advantageous restrictions can be found in the alignments themselves, as researchers often provide a “minimum acceptable” alignment score in the form of a minimum percent identity or maximum number of mismatches. Further, alignments other than the best alignment(s) may be suppressed and pruned from the search space for many use-cases. Certain combinations of these restrictions naturally imply others; for instance, even when no minimum acceptable score is provided, the program can internally track the maximum score obtained for a given query sequence as more alignments are seen and can use that information to ignore reference sequences that have provably sub-optimal matches. This is useful if the program is capable of disqualifying alignments known to be worse than the current best without needing to calculate the entire alignment. Alternatively, if the user specifies a threshold and requires all alignments be reported above it regardless of which are best, the program can still disqualify some alignments based on the provided minimum score.

We describe here an implementation of exhaustive gapped DNA alignment that takes advantage of each of the aforementioned restrictions, together with several additional optimizations discussed in later sections, while guaranteeing that the resulting alignments reported will be identical to those produced by a full exhaustive alignment of all queries against all full-length references. This guarantee holds even in the presence of ambiguous bases (which, except for bases ‘X’ and ‘N’, default to allowing unpenalized matches). We find that our method, BURST, leveraging numerous synergistic optimizations, enables alignment speeds that are up to 20,000 times faster than previously published optimal gapped aligners.

## Results

Here we present BURST, a metagenome-scale DNA alignment software tool which performs exhaustive, optimal alignment to map short reads to a genomic database, guaranteeing the recovery of any or all matches of query sequences to references in the database above a user-specified alignment similarity score (percent identity). A schematic is presented in Figure 1. Various output modes exist with similar runtime performance to output all tied best matches, select a single best match among ties that minimizes the number of unique references chosen, and assign interpolated taxonomy or other semicolon-delimited hierarchical metadata that satisfies a user-specified confidence threshold.

**Figure 1.**
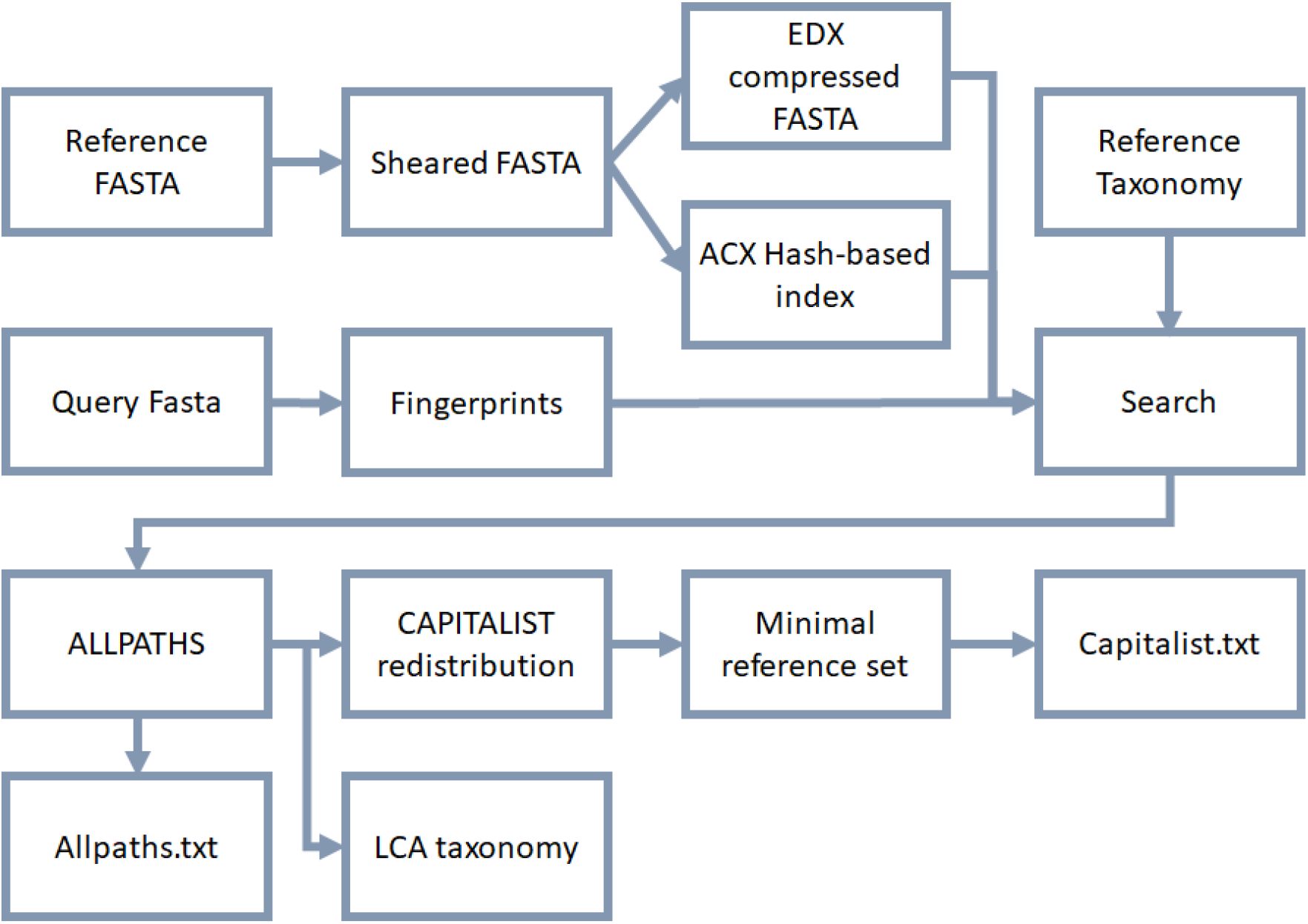
Schematic of BURST. This schematic depicts both the database-generation and alignment arms of BURST. In a common workflow, the user begins by creating a database out of a FASTA of reference sequences, then using that database to align a FASTA of queries and produce alignments with taxonomy according to the desired reporting mode.

To accomplish this with feasible runtime performance, a series of optimizations have been combined as shown in Table 1. It is important to note that only optimizations that do not reduce sensitivity of specificity of alignment are included; they increase speed at no cost of accuracy.

**Table 1.** Overview of algorithmic optimizations in BURST, including estimates of speedup provided by each optimization. BURST uses multiple optimizations to achieve speedup without affecting alignment quality. “+” indicates further speedup is possible in tandem with other items.

| Optimization | Speedup |
| --- | --- |
| **Error-aware pruning** | >2x<sup>1</sup> |
| **Lexicographic table caching** | >2x<sup>1</sup> |
| **Eliminate matrix tracebacks** | 2x |
| **Vectorization along reference clusters using SSE4.1 instructions** | 10x |
| **2-pass search: edit distance then BLAST identity** | 2x |
| **Code micro-optimizations** | 2x |
| **Multithreading: thread-local data with global sync** | +0.9x/thread |
| **Optional high-identity map filter ("accelerator")** | 2-20x<sup>2</sup> |
| **Optional database support with low-identity fingerprint filter** | 1-5x<sup>3</sup> |
| **Lossless prioritization: Ordering, pre-pass, best-hit prioritizing** | 1-10x<sup>3</sup> |

<sup>1</sup> Up to 100x when both caching and pruning are used together
<sup>2</sup> Lower estimate for amplicon databases; higher for shotgun databases
<sup>3</sup> Depends on read length, shearing factor, database uniqueness, and desired identity cutoff

### Performance and system requirements

For full speed, BURST relies on preprocessing of the reference database to produce an optional auxiliary “accelerator” database. This accelerator allows BURST to filter out non-optimal reference sequences, without sacrifice of global optimality, based on patterns of k-mers present in each query sequence and reference sequence. Generally, the memory and disk requirements of BURST are low with only the primary database, but high when an additional accelerator database is produced. The primary database alone occupies space and memory of 0.5-1x the size of the original FASTA file of references. The accelerator, if produced, further occupies ~1-7x the size of the original FASTA file of references in RAM and disk space.

BURST transparently distinguishes between two types of accelerator formats, “small” and “large.” Large format accelerators are produced when the number of reference sequences after shearing and deduplication exceeds 1 million. The RAM and space requirements of both the database and accelerator tends to be just over 7x the size of the original FASTA file of references used to produce them, assuming little redundancy in the original sequences. An example is the “reference and representative genome collection” from NCBI RefSeq(15) v82, which contains just over 5000 representative prokaryotic genomes and very little redundancy. With increasing redundancy, this factor can drop considerably. For example, the space requirement drops to 2.5x the input reference data with over 51,000 prokaryotic genomes, such as those drawn from NCBI RefSeq v84 (all complete, chromosome-, and scaffold-level genomes plus all “representative” contig-level genomes). This factor is expected to drop further with the addition of more genomes as more redundancy is included. For instance, a BURST database of 7,676 *Staphylococcus aureus* genomes from RefSeq v99 (see the GitHub link for the specific accessions) resulted in a database that was less than 1.1x the size of the input FASTA, which is actually smaller than a bowtie2 database of the same (1.9x the size of the input FASTA). Importantly, this bespeaks the increasing scalability of BURST database size as more genomes within a clade become available.

Importantly, BURST demonstrates dramatic speedup over existing optimal algorithms in terms of alignment speed on a real gut amplicon dataset (Table 2). In practical metagenomic alignment, BURST can align, disambiguate, and assign taxonomy at a rate of 1,000,000 150-base query sequences per minute against the RefSeq(15) v82 representative prokaryotic genome database (5,500 microbial genomes, 19GB) at 98% identity on a 32-core computer.

**Table 2.** Alignment through the ages. BURST, an exact method, shows runtime performance comparable to heuristic algorithms on human gut amplicon data using the Greengenes 97% OTU database.

| Year | Method (vs GG97) | Runtime (15M 225bp) |
| --- | --- | --- |
| 1970 | Needleman-Wunsch | 17 years |
| 1999 | GLSEARCH | 310 days |
| 2012 | Bowtie2 (heuristic) | 30 minutes |
| **2017** | **BURST** | **20 minutes** |

**Amplicon test against GreenGenes 13.8 comparing optimal aligners.** The first two rows extrapolated from smaller tests. *Needleman-Wunsch/Smith-Waterman* via “needle”/“water” software *GLSEARCH/SSEARCH* via the FASTA package *Bowtie2* –very-sensitive included as (heuristic) speed anchor

A comparison of BURST database creation times is also presented in Table 3. Both aligners were run with 28 threads on a dual Xeon Platinum 8180 system with 1.5TB RAM.

**Table 3.** A comparison of BURST database creation times. The *S. aureus* database is as described previously and available on the project GitHub. The “Rep 200” dataset is composed of RefSeq v200 assemblies marked as “representative” or “reference” genomes in RefSeq assembly_summary manifest.

| Aligner | DB creation time (m) | RAM use (GB) | Size of index (GB) |
| --- | --- | --- | --- |
| BURST v1.0 (*S. aureus*) | 106 | 183 | 22 |
| Bowtie2 v2.5.3.1 (*S. aureus*) | 714 | 68 | 39 |
| BURST v1.0 (Rep 200) | 83 | 500 | 166 |
| Bowtie2 v2.5.3.1 (Rep 200) | 632 | 141 | 95 |

### Availability

BURST is available under the AGPLv3 license. Precompiled static binaries (with no dependencies or installation required other than simply downloading and running BURST). It is available on GitHub at github.com/knights-lab/BURST (see the releases page under “Downloads” for the precompiled binaries: github.com/knights-lab/BURST/releases).

## Methods

### Alignment

The alignment scoring function proposed and used in this work is fundamentally a dynamic-programming-based similarity measure similar to the Needleman-Wunsch algorithm (3) with a two-component objective function, the first of which is Levenshtein distance(16) and the second being alignment identity (sometimes called “BLAST id”). Creation of this hybrid metric was informed by a desire to score internal alignments using criteria consistent with those commonly used later to filter them. The advantage in using the hybrid measure over either component alone is that the Levenshtein distance alone is subject to producing multiple different but identically-scoring alignments from a single short-read alignment, and the BLAST id may produce a slightly higher-scoring alignment that is substantially longer and more gap-ridden, and hence of dubious biological merit. Using BLAST id to pick from among those alignments produced by Levenshtein distance provides a natural resolution. Another commonly used metric for screening and evaluating alignments, the e-value, was not considered as BURST performs end-to-end alignments only, and in context of short-read mapping, the query sequences are often of similar or identical lengths as one another, reducing the discriminative utility of the e-value in this context.

Although the omission of alternative scoring criteria such as variable gap penalties or nucleotide-specific scores may be unreasonable for general-purpose alignments(17,18), their utility in the high-identity shortread mapping problem is lower. The objective of short-read mapping, rather than characterizing local evolutionary homology as with comparison of two full-length protein sequences, is instead tantamount to fuzzy substring search where the relatively few allowable mismatches can be contributed by artifactual sequencing noise or recent evolutionary divergence.

### “Synergistic” caching and pruning

Within the alignment matrix, two major optimizations are used which in tandem increase the rate of alignment more than either alone. The first such optimization is lexicographic caching of previous query alignments to the same reference, eliminating the need to recalculate this matrix for a number of rows equal to the length of the shared prefix. Importantly, for such an optimization to be useful, query sequences are sorted using a fast multi-threaded sorting method to maximize the length of the shared prefix between adjacent query sequences. For 1 million random sequences with 4-letter alphabet (DNA), it is easy to observe that the length of a shared prefix of length 10 will be observed, on average, 1-(1-(1/4)^10^)^1000000^ or 61.5% of the time between any two adjacent queries. For amplicon data, the shared prefix length is typically much higher as the selection of genomic region in this case is non-random. Figure 2 (blue cells) shows an example of this optimization (we can assume the previous query sequence is “TCGAACAA”).

**Figure 2.**
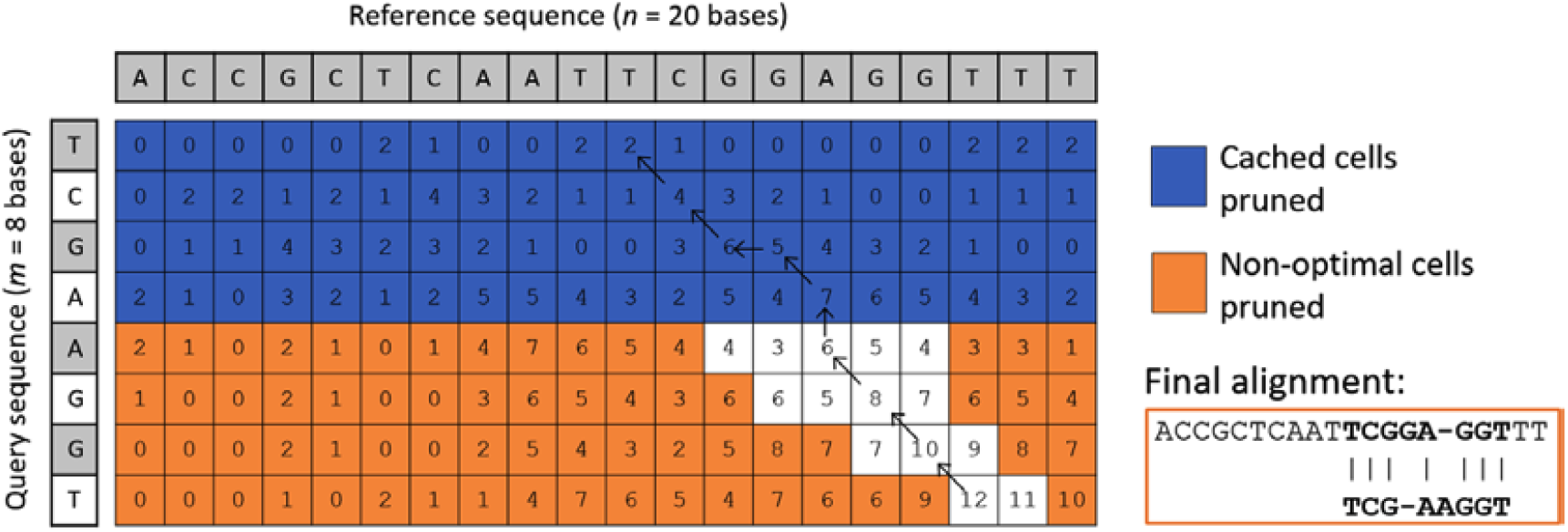
Simplified example DNA alignment accelerated by BURST. Optimal alignments usually require filling a dynamic programming matrix from top-left to bottom-right and finding the highest-scoring path. BURST skips calculating all blue and orange cells while guaranteeing the best alignment (saving 146/160 = 91.25% of the computation). The previous query’s alignment (not shown) is cached until the point of prefix divergence from the current (blue). Also, cells that cannot form future valid alignments given current scores (orange) are not pursued. Path traceback (arrows) is also eliminated.

The key synergistic optimization used in tandem with this caching step is opportunistic pruning of guaranteed “dead end” paths through the dynamic programming matrix (Figure 2, orange cells). For each cell, a threshold score calculated by summing that cell’s current alignment score with the maximum possible score for all remaining nucleotides in the query, assuming all receive perfect matches from this query position onward. The cell is marked inactive if this threshold score is lower than the minimum user-defined score (or the best score among all previous alignments of this query, if in a “best-hit” reporting mode). Elimination of traceback is also implemented so as not to require final traversal through the best-scoring path at the end of alignment; this is accomplished by storing an auxiliary matrix that tracks gap counts directly rather than noting the direction of the best-scoring path at each cell (Figure 2, black arrows).

Another strategy for improving runtime performance is the use of a 2-pass scoring criterion for determination of best matches. Because the Levenshtein distance calculation is prerequisite for the secondary BLAST id selection, the former alone can be applied as an initial screening pass; if an alignment fails the first criterion, it will by definition fail the second and be rejected, sparing the need to compute both for each alignment.

### BURST enables arbitrarily confident taxonomy assignment per read with thresholded LCA

In one of its modes of operation, BURST can assign taxonomy (or any other hierarchical, semicolon-delimited feature) to each read using lowest common ancestor (LCA).

## Discussion

BURST is a high-performance standalone program with no dependencies that enables exhaustive, high-throughput metagenomic alignment without heuristics or approximations. This makes BURST uniquely suitable for a variety of tasks where accuracy is of paramount importance, such as part of a pathogen screening or strain detection pipeline. As shown in Figure 3a, the heuristic aligner bowtie2(9) increasingly fails to discover optimal matches with higher sequence divergence from known references, whereas BURST’s best hit always returns an optimal match. Not only does BURST provide more accurate best hits with decreasing sequence similarity, it also enables some novel use-cases as well.

**Figure 3.**
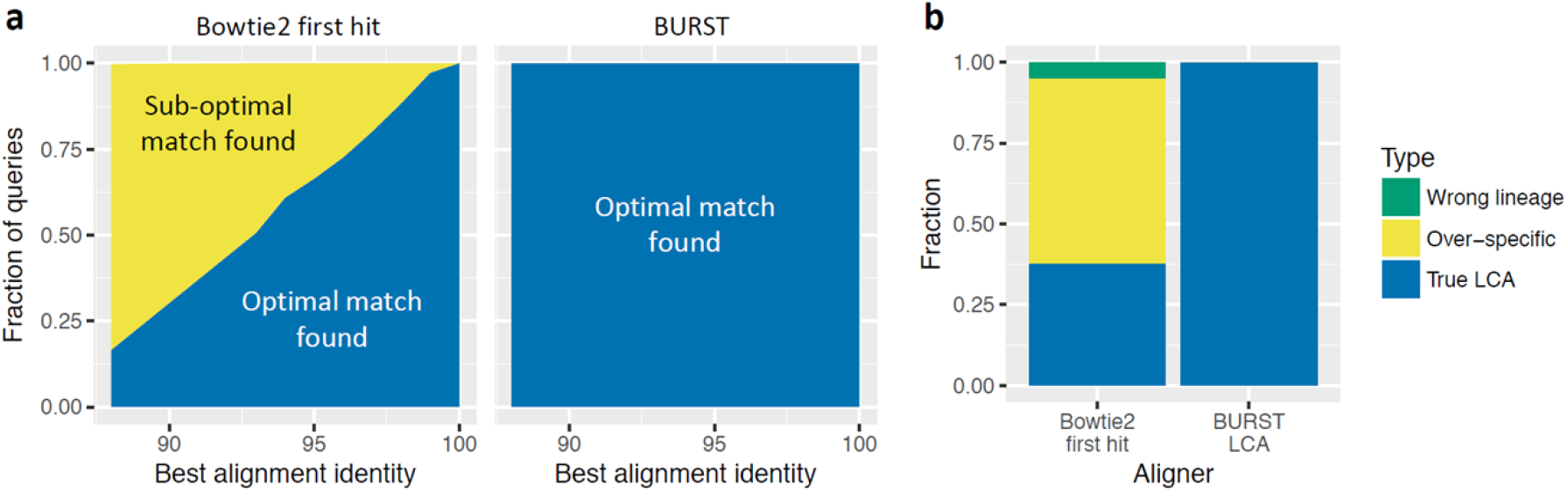
BURST vs Bowtie2: best hit and LCA taxonomy (a) A comparison of BURST in best-hit mode compared to the default runtime configuration of bowtie2 on an example gut amplicon dataset against the Greengenes 97% OTU database. BURST recovers an optimal best-scoring match for all queries at all alignment similarities, while bowtie2 returns more suboptimal matches with decreasing alignment identity. (b) A comparison of BURST in LCA mode vs bowtie2 default settings in terms of reported taxonomy. When multiple identically-scoring best hits exist, BURST will return the lowest common ancestor (LCA) as the taxonomy, while best-hit methods pick an arbitrary representative which is often over-specific or, due to heuristics, sometimes the wrong lineage entirely.

BURST can be used to report conservative taxonomy on a per-read basis by finding the lowest common ancestor among all tied alignments against all references for each query. Using this approach on an example human amplicon dataset(19) with the Greengenes(20) 13_8 reference database, we find that using a naïve “best-hit” approach for assigning taxonomy results in drastic levels of overconfidence in taxonomy assignments (Figure 3b). This overconfidence can lead to split signals when, for instance, multiple different species of a genus are reported when they do not actually exist in a sample. In this case, BURST would report higher amounts of the genus alone without assigning arbitrary species labels at random.

When used in “Forage” mode, BURST can also be used as an exhaustive means to determine where a set of primers may align in a database of many genomes, including the ability to specify primers with ambiguous bases and any number of mismatches, insertions, or deletions. Additionally, using BURST’s ability to report all tied best alignments (“ALLPATHS” mode) can be used in interesting ways such as in Figure 4, which shows a Cytoscape visualization of the relatedness of 21 strains within three species of the bacterium *Enterococcus*. For *E. faecalis* and *E. faecium*, 10 strains each were randomly selected from RefSeq(15) v86 assemblies, and a single strain of *E. aquimarinus* was used. As expected from a phylogenetic analysis(21), *E. faecalis* and *E. faecium* are substantially more closely related than either is to *E. aquimarinus*. Ecologically, the former two microbes are also common gut commensals, whereas *E. aquimarinus* was isolated from sea water(22). It may also be inferred that the randomly selected E. faecalis strains are more dissimilar to one another than the E. faecium strains. This may have important implications for creation of a reference database, as intelligent strain selection may be beneficial to capture a species’ full pan-genomic content without inflating the size of the database with highly similar strains. More fundamentally, it visually highlights the fact that not all species of a genus are equidistant from one another, nor are strains of a species.

**Figure 4.**
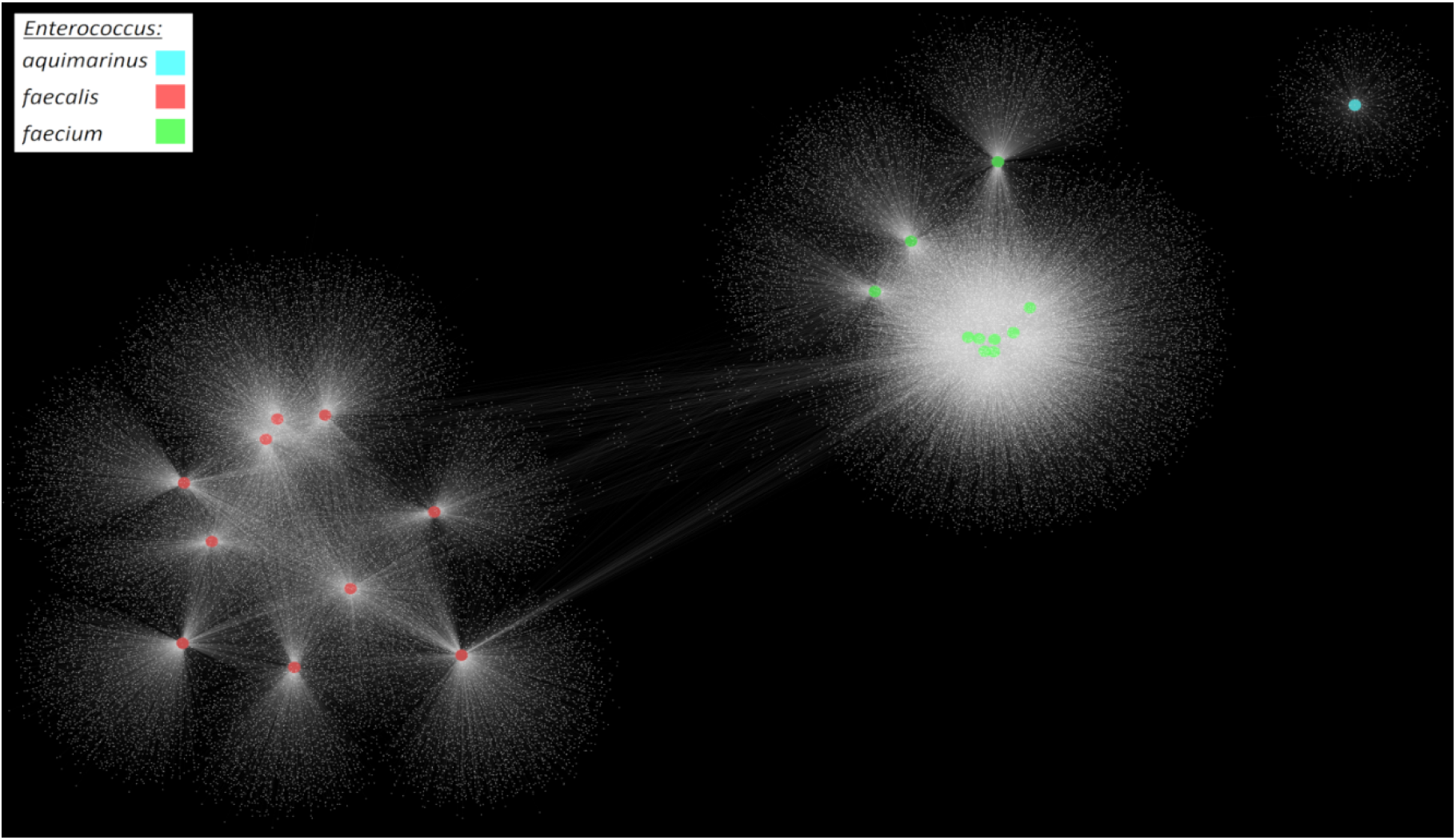
BURST reveals species relatedness through shared genes. All genes from 21 strains of Enterococcus were aligned with BURST in “ALLPATHS” mode against a database of the same strains’ genomes. Large nodes indicate individual genomes, colored by species, and white dots represent individual genes. Lines indicate alignments between genes and genomes. Force-directed layout was used to automatically orient the network. The randomly selected *E. faecalis* and *E. faecium* share more genes with themselves than with each other, and the *E. aquimarinus* shares little with either other species.

Another reporting mode in BURST that may be especially useful for whole-genome metagenomics is the “capitalist” reporting mode, a Minpath-like(23) parsimony-based disambiguation scheme the workings of which are highlighted in Figure 5a in comparison to the conservative LCA method (Figure 5b) also demonstrated above. An example of running this mode on a simulated dataset is shown in Figure 6.

**Figure 5.**
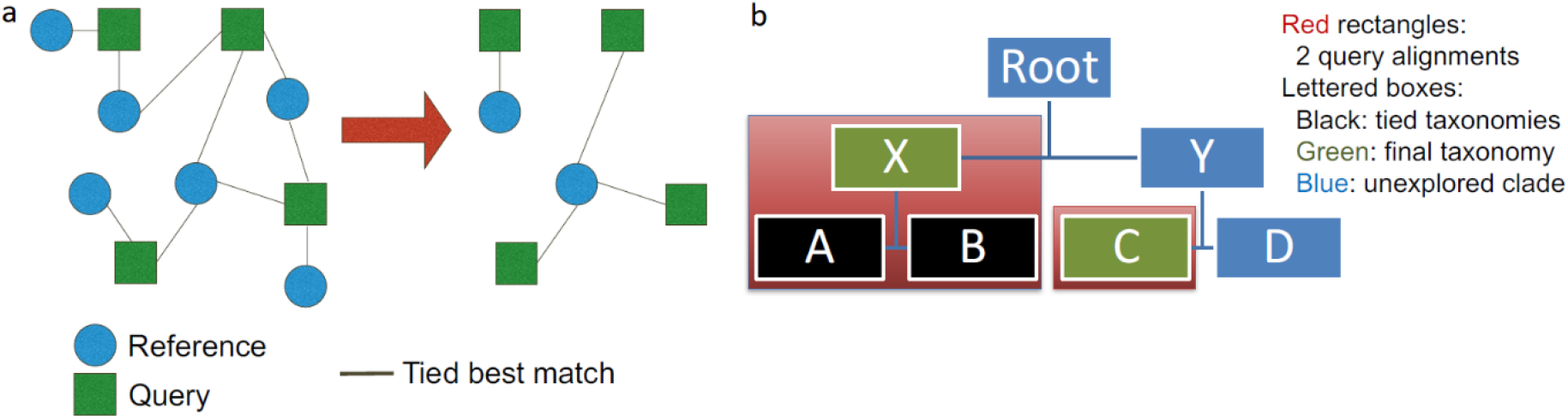
Illustration of BURST “Capitalist” disambiguation vs LCA. (a) Tied best matches arise due to repetition in multiorganism reference databases. To resolve this, BURST computes the minimal set of references necessary to satisfy all queries. This reduces split statistical signals present in most metagenomic data. (b) Lowest common ancestor (LCA) is an alternative, more conservative approach that independently assigns each read a taxonomy by backtracking from leaf to root (most specific to least specific). BURST implements both methods.

**Figure 6.**
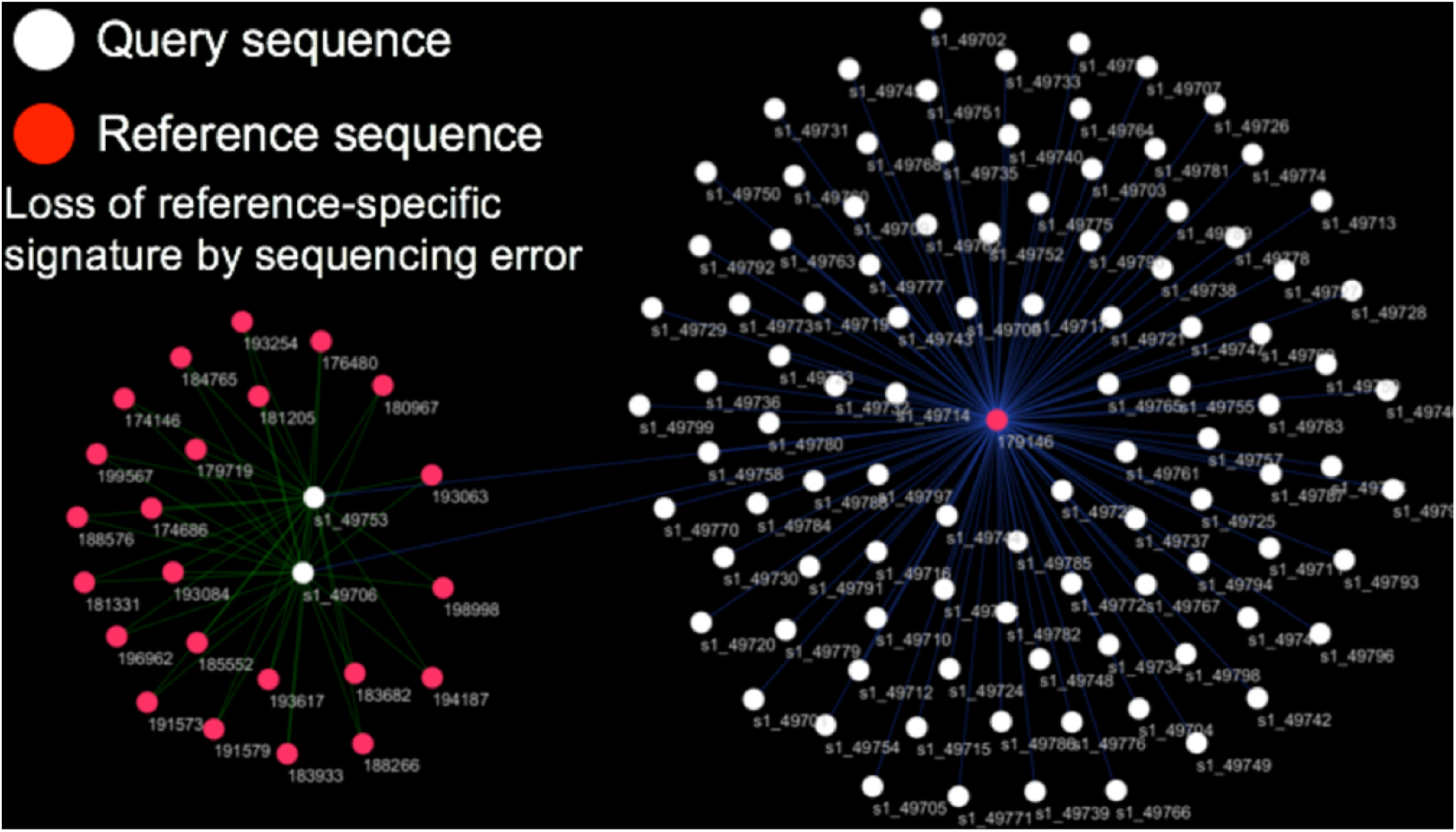
Practical example of CAPITALIST. In this example run on a simulated dataset, all of the queries (white dots) originate from the same reference (red dot in right-center). However, in two of these queries (two white dots on the left-center), the nucleotides necessary to confer unambiguous best alignments to their reference of origin were mutated, making them align equally well to a number of other references as well. Lines represent all possible tied best alignments; blue lines represent capitalist picks (minimizing alignments), which would eliminate all extraneous references on the left and properly assign the two left-most queries to the same reference as the other queries.

## Conclusion

With its favorable performance attributes and guarantee of alignment optimality, BURST may be a reasonable alternative to other aligners for metagenomic and genomic DNA alignment. Further, BURST incorporates a number of features useful for metagenomic analysis including methods to disambiguate ties, assign conservative taxonomy annotations, and reduce database size with increasing numbers of similar reference genomes. Because it is not splice-aware, nor does it support human-centric alignment formats such as SAM(24), it may not be immediately suitable for human genomic applications reliant on these features. Although BURST is up to 20,000 times faster than previous optimal gapped alignment algorithms, it will still be 10-to-100 times slower than some heuristic or non-optimal search methods such as k-mer-based search. For whole-genome databases, BURST also relies heavily on a large precalculated database accelerator file that can reach upwards of 7x the input (FASTA) database size. Therefore, researchers may not wish to use BURST if speed and low-RAM usage are more important than optimality.

Further improvements to BURST may include the ability to align protein sequences (including local alignments within a specified minimum length and support for distance matrices) as well as further improvements to the compressive database engine and other optimizations to increase speed and reduce memory usage. Improvements to the reporting of alignments may also include incorporation of a “coverage”-based statistic, the goal of which would be to disambiguate reads based not only on aggregate parsimony but also on evenness of read coverage throughout the genome. Future work may characterize the extent to which the various output modes of BURST impact the performance of statistical and machine learning models.

